# Emergence of antibiotic-specific *Mycobacterium tuberculosis* phenotypes during prolonged treatment of mice

**DOI:** 10.1101/2024.08.20.607990

**Authors:** Elizabeth A Wynn, Christian Dide-Agossou, Reem Al Mubarak, Karen Rossmassler, Victoria Ektnitphong, Allison A Bauman, Lisa M Massoudi, Martin I Voskuil, Gregory T Robertson, Camille M Moore, Nicholas D Walter

**Author notes:** Nicholas D Walter and Camille M Moore contributed equally to this work. Corresponding author: Nicholas D Walter.

## Abstract

A major challenge in tuberculosis (TB) therapeutics is that antibiotic exposure leads to changes in the physiologic state of *M. tuberculosis* (*Mtb*) which may enable the pathogen to withstand treatment. While antibiotic-treated *Mtb* have been evaluated in short-term *in vitro* experiments, it is unclear if and how long-term *in vivo* treatment with diverse antibiotics with varying treatment-shortening activity (sterilizing activity) affect *Mtb* physiologic states differently. Here, we used SEARCH-TB, a pathogen-targeted RNA-sequencing platform, to characterize the *Mtb* transcriptome in the BALB/c high-dose aerosol infection mouse model following 4-week treatment with three sterilizing and three non-sterilizing antibiotics. Certain transcriptional changes were concordant among most antibiotics, including decreased expression of genes associated with protein synthesis and metabolism, and the induction of certain genes associated with stress responses. However, the magnitude of this concordant response differed between antibiotics. Sterilizing antibiotics rifampin, pyrazinamide, and bedaquiline generated a more quiescent *Mtb* state than did non-sterilizing antibiotics isoniazid, ethambutol, and streptomycin, as indicated by decreased expression of genes associated with translation, transcription, secretion of immunogenic proteins, metabolism, and cell wall synthesis. Additionally, we identified distinguishing transcriptional effects specific to each antibiotic, indicating that different mechanisms of action induce distinct patterns of cellular injury. In addition to elucidating *Mtb* physiologic changes associated with antibiotic stress, this study demonstrates the value of SEARCH-TB as a highly granular pharmacodynamic assay that reveals antibiotic effects that are not apparent based on culture alone.

## INTRODUCTION

Tuberculosis (TB) is the leading cause of death from infection globally, killing approximately 1.2 million people each year.^1^ Because standard antibiotic treatment regimens require 4 to 6 months to reliably cure drug-susceptible TB,^2^ there is an urgent need for new antibiotic combinations capable of curing all forms of TB more quickly.^3^

One reason that months-long treatment is required to reliably cure TB is that antibiotic exposure changes the physiologic state of *M. tuberculosis* (*Mtb*).^4^ The physiologic state of *Mtb* is a key determinant of antibiotic activity.^5–8^ However, there is a paucity of information about the physiologic processes of *Mtb* in an *in vivo* setting or how they might differ depending on an antibiotic’s mechanism of action. Attention has historically focused on the direct mechanism of action of antibiotics (*i.e.,* the molecular interaction of an antibiotic with its target protein). However, for *Mtb* that are not immediately killed by initial antibiotic exposure, the immediate injury caused by antibiotic-target binding initiates a cascade of secondary, indirect physiologic perturbations,^9^ resulting in chronically stressed bacteria. *Mtb* that survive long-term treatment, and therefore have the potential to cause relapse, are likely shaped by the specific nature of the injury (*i.e.,* the mechanism of action of a given antibiotic). The effect of antibiotics on *Mtb* physiologic processes has been studied extensively in short-term *in vitro* experiments,^10–15^ but short-term exposure in axenic culture may not replicate the physicochemical conditions and dynamic pharmacokinetics encountered during chronic *in vivo* exposure. Here, the use of a novel targeted RNA-seq platform called SEARCH-TB^16^ enabled us to characterize *Mtb* that emerge during prolonged treatment with diverse antibiotics in mice.

While all antibiotics included in conventional combination regimens are thought to contribute to cure to some degree, certain antibiotics play a more pronounced role in shortening the time required to cure TB.^17^ Historically, antibiotics such as rifampin, pyrazinamide, and bedaquiline, which have potent treatment-shortening activity, have been described as “sterilizing” while antibiotics such as isoniazid, streptomycin, and ethambutol, which may have bactericidal activity but contribute only modestly to shortening the time needed to achieve a non-relapsing cure, have been described as “non-sterilizing.”^18^

In this study, we compared the long-term effect of three canonical sterilizing antibiotics (rifampin, bedaquiline, pyrazinamide) and three canonical non-sterilizing antibiotics (isoniazid, streptomycin, ethambutol) over a 28-day treatment period in the BALB/c high-dose aerosol infection mouse model. We first identified *Mtb* transcriptional changes that were common to most of the antibiotics assessed, then compared the effect of sterilizing versus non-sterilizing antibiotics, and finally characterized transcriptional features unique to each antibiotic.

## METHODS

### 1. Murine experiments and RNA extraction

Experiments used the BALB/c high-dose aerosol infection model, which is central to contemporary TB drug development.^19^ Female BALB/c mice, 6 to 8 weeks old, were exposed to aerosol (Glas-Col) with *Mtb* Erdman strain, resulting in 4.55 ± 0.03 (SEM) log_10_ colony forming units (CFU) in lungs on day one. Mice euthanized after 11 and 19 days (when clinical deterioration required euthanasia) served as pre-treatment and untreated control groups. Starting day 11, mice were treated via oral gavage five days a week for 28 days before euthanasia. We used established human-equivalent doses of all antibiotics (Table S1) except for bedaquiline which was administered at one-fifth of the human-equivalent dose because the full human-equivalent dose resulted in *Mtb* burden too low for reliable SEARCH-TB profiling. Lungs were flash frozen before CFU enumeration and RNA extraction as recently described.^16^ All animal procedures were supervised by the Colorado State University Animal Care and Use Committee and conducted according to established guidelines.

### 2. RNA sequencing, and data preparation

Sequence analysis of samples was performed via SEARCH-TB following recently described methods.^16^ Briefly, RNA was reverse transcribed, and cDNA targets were then amplified using the SEARCH-TB panel. Libraries were sequenced on an Illumina NovaSeq6000. We followed the bioinformatic analysis and quality control pipeline as recently described.^16^

### 3. Statistical Analysis

Following normalization with DESeq2’s variance stabilizing transformation (VST),^20^ we performed principal component analysis (PCA) on the 500 most variable genes. We estimated differential expression by fitting negative binomial generalized linear models to each gene using edgeR.^21^ Likelihood ratio tests were used to compare gene expression between groups.

To identify groups of genes with similar expression patterns across conditions, we performed hierarchical clustering of the predicted expression values obtained from the edgeR models after filtering out invariant genes (*i.e.,* not differentially expressed between any two conditions). Then, using Euclidean distance with Ward’s method,^22^ we clustered the genes based on the predicted expression values for each condition. To further visualize the expression patterns for individual clusters, we used sample-specific, scaled VST normalized expression values averaged across the genes in each cluster (Figure S1). Using analysis of variance (ANOVA) and post-hoc pairwise t-tests, we evaluated between-group differences for each cluster using these scaled expression values.

We performed functional enrichment analysis using gene categories established by Cole et al.^23^ and curated from the literature (Table S2) using the hypergeometric test in the hypeR package^24^ to evaluate whether genes differentially expressed in pairwise comparisons between conditions were overrepresented in each gene set. Enrichment analysis was run twice, using significantly upregulated and significantly downregulated genes separately. Gene categories with fewer than 8 genes were excluded. All analyses were performed using R (v4.3.1) and comparisons were considered statistically significant when Benjamini-Hochberg adjusted *p*-values^25^ were less than 0.05. Gene expression for individual gene categories was visualized using sample-specific scaled VST normalized expression values averaged across the genes in the category (Figure S1). Differential expression, functional enrichment, and visualizations can be evaluated interactively using an Online Analysis Tool [https://microbialmetrics.org/analysis-tools/].

## RESULTS

### 1. Bactericidal effect of antibiotic treatments

We first characterized the antibiotic effect based on changes in colony forming units (CFU), which estimates the number of bacilli capable of growth on solid agar (**Fig. 1a**). In pre-treatment control mice sacrificed on post-infection day 11, the average lung CFU burden was

**Figure 1.**
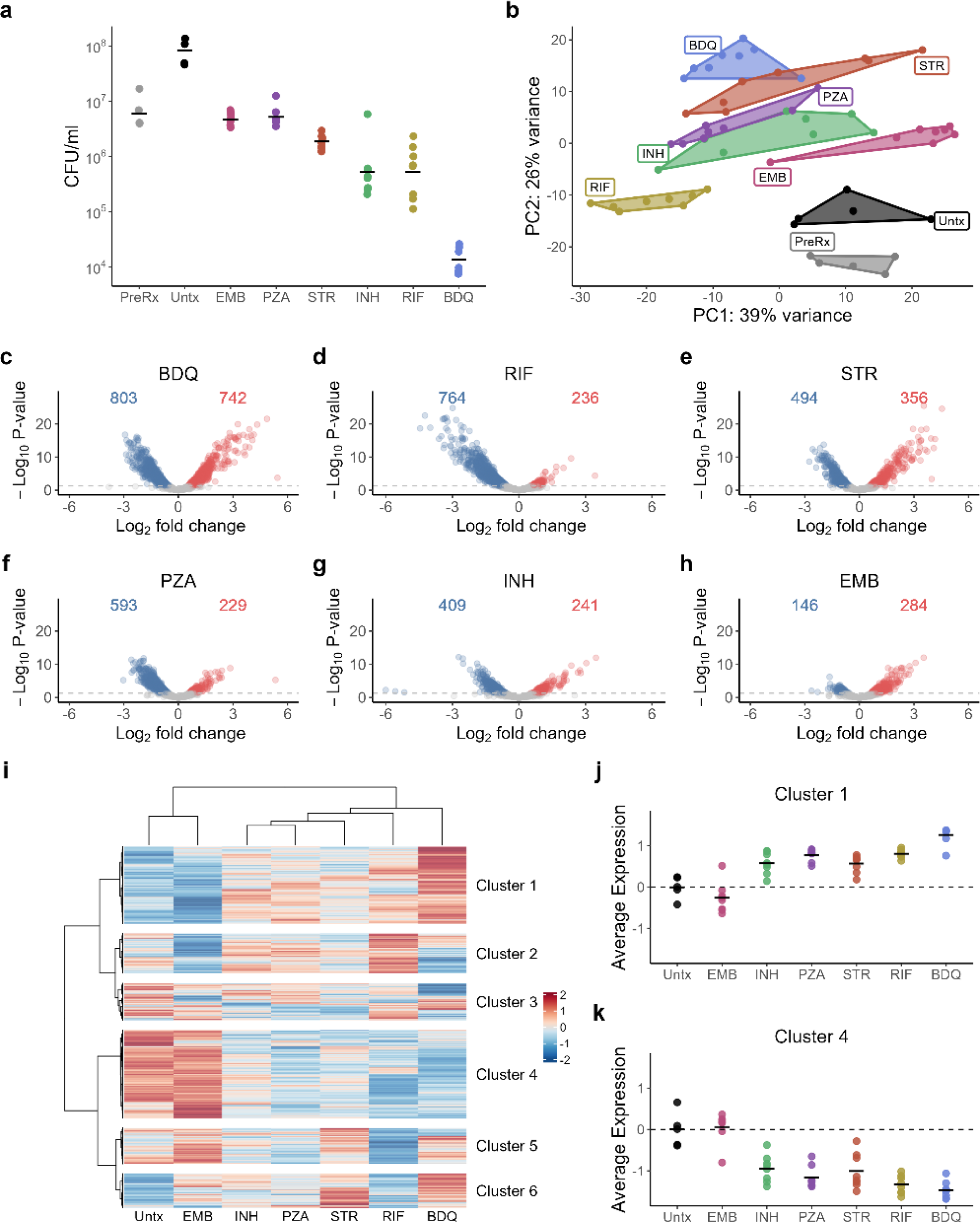
**a**. *Mtb* CFU burden in the lungs of BALB/c mice after 4-week treatment with individual antibiotics. Circles indicate CFU values from individual mice. Horizontal bars indicate group means. PreRx and Untx indicate control mice sacrificed on the day treatment was initiated or 8 days thereafter, respectively. **b**. Principal Components Analysis (PCA) plot of VST-normalized gene expression data, for the top 500 most variable genes. The first two principal components are shown on the x- and y-axes and each point represents an individual sample. A convex hull highlights antibiotic treatments. **c-h**. Volcano plots summarizing the differential expression between *Mtb* in untreated mice and *Mtb* in (c) BDQ, (d) RIF, (e) STR, (f) PZA, (g) INH, and (h) EMB. The number of genes significantly down- (blue) or upregulated (red) for each antibiotic treatment relative to untreated (adj *p*-value< 0.05) are shown. **i.** Heatmap of gene expression including all genes significantly differentially expressed between at least two treatment conditions (N=2,589). Values are row-scaled, with red and blue indicating higher and lower expression, respectively. Hierarchical clustering of genes identified six broad patterns. **j-k**. Average of VST-normalized, scaled expression across treatments for clusters (j) one and (k) four. Each point represents an individual mouse. Horizontal lines indicate average values. Values are centered around the average value for the untreated samples so that points above and below zero represent upregulation or downregulation relative to untreated, respectively. Abbreviations: Untreated (Untx), Ethambutol (EMB), Isoniazid (INH), Pyrazinamide (PZA), Streptomycin (STR), Rifampin (RIF), Bedaquiline (BDQ).

6.78 log_10_. In untreated control mice, which were maintained without treatment until post-infection day 19 when clinical deterioration required euthanasia, the average lung CFU burden was 7.91 log_10_. The average increase of 0.14 log_10_ per day between days 11 and 19 indicated rapid bacterial replication. Pyrazinamide and ethambutol had a static effect, preventing an increase in CFU burden relative to the pre-treatment control but not reducing the CFU burden after 28 days of treatment. Streptomycin reduced lung CFU by 0.5 log_10_ relative to the pre-treatment control. Rifampin and isoniazid had bactericidal activity, reducing CFU by 1.05 and 1.06 log_10_ relative to pre-treatment control, respectively. Bedaquiline had the greatest bactericidal effect, reducing CFU by 2.64 log_10_.

### 2. Clustering of antibiotic-induced transcriptional change

Principal Component Analysis of the SEARCH-TB results showed that samples from each antibiotic clustered distinctly from one another (**Fig. 1b**), demonstrating that antibiotics with unique mechanisms of action affect *Mtb* differently. The untreated control (19 days after aerosol infection) was distinct from the pre-treatment control (11 days after aerosol infection), consistent with the effect of adaptive immunity, which is known to occur around day 14.^26^ To isolate the effect of antibiotics rather than immunity, we selected the untreated control as our primary reference. The number of *Mtb* genes significantly altered by antibiotic exposure ranged from 430 (ethambutol) to 1,545 (bedaquiline) (**Fig. 1c-h**), indicating that each antibiotic stress induced broad changes in bacterial physiological state. To visualize the differences between antibiotics, we performed unsupervised hierarchical clustering based on the average expression of differentially expressed genes (**Fig. 1i**). Of the antibiotics evaluated, ethambutol was the most similar to the untreated control. Isoniazid, streptomycin, pyrazinamide, and rifampin clustered together and were distinct from the transcriptional changes caused by bedaquiline.

Unsupervised clustering of differentially expressed genes revealed that certain clusters of genes behaved concordantly among most antibiotics, while others behaved discordantly. For all antibiotics except ethambutol, genes in Cluster 1 (N=639) exhibited increased expression relative to untreated control (**Fig. 1i**). The magnitude of induction of Cluster 1 genes varied between antibiotics (**Fig. 1j**), with greater increase for bedaquiline than for any other antibiotic (*p-*value relative to the closest antibiotic= 0.0003). Conversely, for all antibiotics except ethambutol, genes in Cluster 4 (N=731) had decreased expression relative to the untreated control (**Fig. 1i**), with greater decreases for bedaquiline and rifampin than for isoniazid, streptomycin, and pyrazinamide (**Fig. 1k**). The remaining clusters (2, 3, 5, and 6) identified genes affected in distinct ways by antibiotics with different mechanisms of action (average expression plots in Supplemental Fig. S2), consistent with the emergence of antibiotic-specific injury responses that are discussed further below. Functional enrichment for each cluster is summarized in Supplemental File 1.

### 3. Concordant *Mtb* transcriptional responses to diverse antibiotic exposures

This section describes the transcriptional responses that were shared among most antibiotic exposures. As described above, ethambutol did not change CFU, clustered with the untreated control and had the smallest number of differentially expressed genes relative to untreated control (**Fig 1h**). To characterize effective antibiotic treatment, ethambutol was therefore excluded from our description of concordant transcriptional responses below. For individual mice, we summarized the average normalized expression of genes in established biological categories (Supplemental Table S2) (**Fig. 2**). For each of the Figure 2 plots, Supplemental Figure S3 includes a corresponding heatmap that summarizes the average expression of individual genes in each category. Statistical results of the functional enrichment analysis are provided in Supplemental File 2.

**Figure 2.**
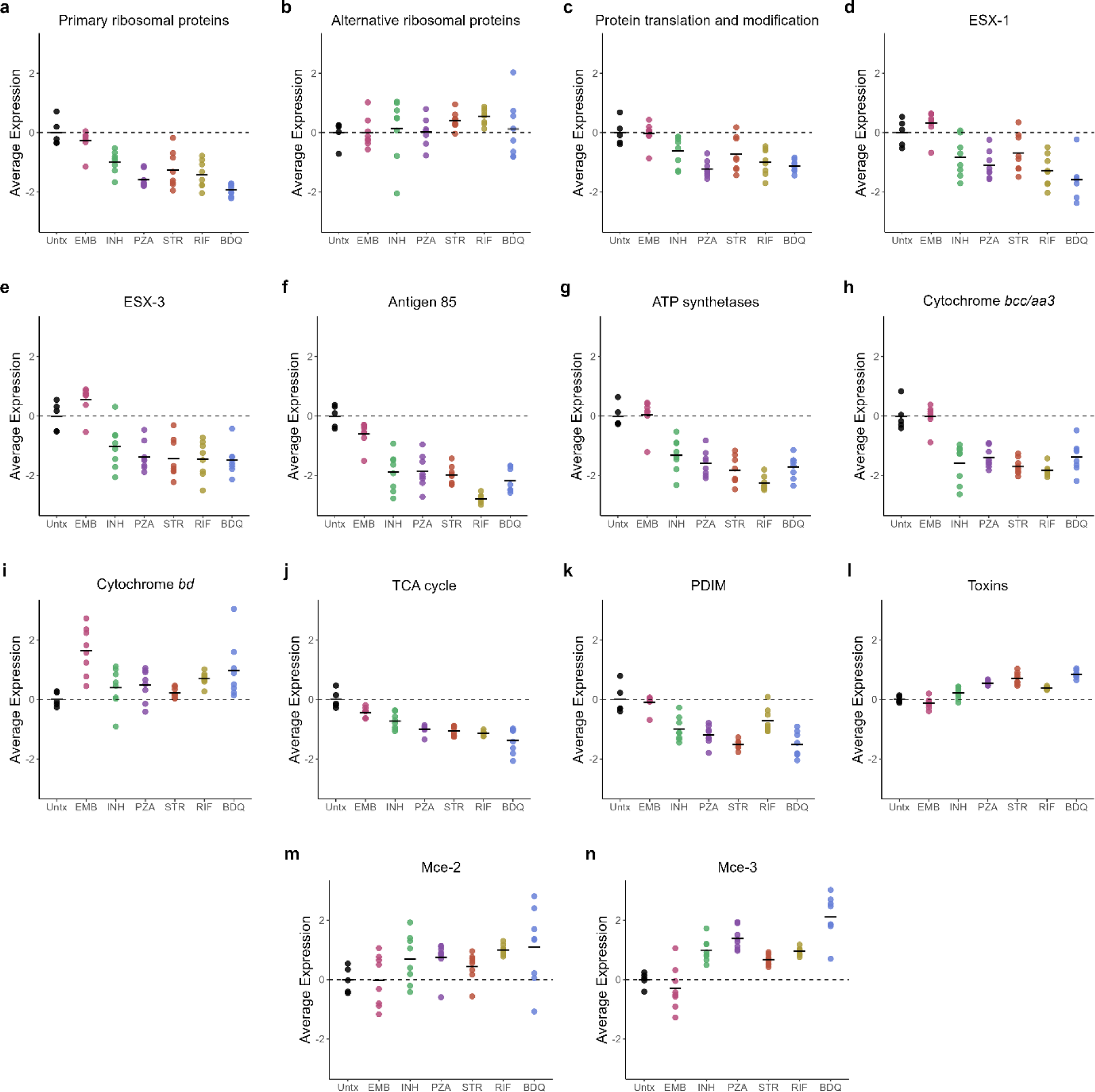
Average of VST-normalized, scaled gene expression across *Mtb* treatments in BALB/c mice for genes in key *Mtb* biological processes: (**a**) primary ribosomal proteins, (**b**) alternative ribosomal proteins, (**c**) protein translation and modification, (**d**) ESX-1, (**e**) ESX-3, (**f**) antigen 85, (**g**) ATP synthesis, (**h**) cytochrome *bcc/aa3*, (**i**) cytochrome *bd*, (**j**) TCA cycle (**k**) PDIM, (**l**) toxins, (**m**) MCE-1, and (**n**) MCE-3. Each point represents an individual mouse. Horizontal lines indicate average values. Values are centered around the average value for the untreated samples so that points above and below zero represent upregulation or downregulation relative to untreated, respectively. Abbreviations: Untreated (Untx), Ethambutol (EMB), Isoniazid (INH), Pyrazinamide (PZA), Streptomycin (STR), Rifampin (RIF), Bedaquiline (BDQ)

#### Suppressed expression of genes associated with protein translation

Antibiotics concordantly decreased the expression of the primary ribosomal protein genes relative to the untreated control, consistent with slowing of protein synthesis (**Fig. 2a**). By contrast, the four “alternative" ribosomal protein genes involved in stress-induced ribosomal remodeling^27,28^ had sustained or increased expression (**Fig. 2b**) (gene set too small for statistical functional enrichment evaluation). Antibiotics decreased expression of the protein translation and modification category that includes genes responsible for translational initiation, promotion of tRNA binding, elongation, termination, and protein folding (**Fig. 2c**) (statistically significant in functional enrichment analysis for pyrazinamide, rifampin, and bedaquiline).

#### Decreased expression of immunogenic secretory proteins

Relative to untreated control, antibiotics decreased expression of the ESX-1 secretion system, including *esxA* and *esxB,* which encode the highly-immunogenic early secretory antigenic 6 kDa (ESAT-6) and culture filtrate protein 10 (CFP-10), respectively (**Fig. 2d)** (statistically significant in functional enrichment analysis for all except ethambutol and streptomycin). Antibiotics decreased expression of the ESX-3 system that secretes peptides that activate neutrophil and macrophages (**Fig. 2e**). Finally, antibiotics appeared to decrease expression of the three genes coding for the Antigen 85 complex (**Fig. 2f**), a secreted protein essential for survival within macrophages which also helps to maintain the *Mtb* cell wall integrity by catalyzing the transfer of mycolic acids to cell wall (gene set too small for statistical functional enrichment analysis).^29^

#### Metabolic slowing and adaptation

Relative to the untreated control, antibiotics significantly suppressed expression of genes coding for ATP synthetases (**Fig. 2g**). Oxidative phosphorylation appeared to transition from the primary cytochrome *bcc/aa3* supercomplex (downregulated) to the less-efficient cytochrome *bd* oxidase (upregulated), which has been implicated in persistence under environmental and antibiotic stress^30^ (**Fig. 2h-i**) (gene sets too small for statistical functional enrichment evaluation). Antibiotics were associated with decreased expression of TCA cycle genes (**Fig. 2j**) (all except ethambutol and rifampin were statistically significant in functional enrichment analysis). Respiratory slowing was not accompanied by the expected increased expression of glyoxylate bypass genes, an alternative pathway previously implicated in antibiotic tolerance.^31^ Genes associated with carbon storage such as triacylglycerol were also not upregulated. Specifically, *tgs1,* a gene in the DosR regulon which codes for triacylglycerol synthase previously associated with lipid accumulation during treatment,^32^ had significantly decreased expression after exposure to all drugs except ethambutol and isoniazid (see Online Analysis Tool).

#### Decreased synthesis of mycolic acids and PDIM

Antibiotics significantly reduced the expression of Rv2524c (*fas*), the gene coding for fatty acid synthetase I, indicating a slowdown in the first step of mycolic acid synthesis (see Online Analysis Tool). All antibiotics except ethambutol appeared to decrease expression of

Phthiocerol dimycocerosate (PDIM), suggesting potential decreased virulence of the antibiotic-stressed phenotypes^33^ (**Fig. 2k**) (statistically significant in functional enrichment analysis for all antibiotics except ethambutol and rifampin).

#### Regulation of growth: sigma factors

Consistent with transition to a quiescent phenotype, antibiotics resulted in significantly lower expression of *sigA*, which codes for the primary ‘housekeeping’ sigma factor necessary for growth, relative to untreated control (see Online Analysis Tool). Other sigma factors were affected differently by individual antibiotics and are discussed in Section 5 below.

#### Modulation of stress responses

Antibiotics induced expression of genes for toxins that act post-transcriptionally to reprogram *Mtb* in response to stress (**Fig. 2l**) (statistically significant in functional enrichment analysis for streptomycin, pyrazinamide, and bedaquiline). However, as described below, the pattern of which toxin genes had increased expression differed depending on antibiotic exposure. Consistent with the change previously observed with the standard 4-drug regimen,^16^ mammalian cell entry (*mce*) operons, initially identified as *Mtb* virulence adaptations and more recently implicated in stress adaptation,^34^ appeared to have increased expression of Mce-2 and Mce-3 operons with all drugs except ethambutol (**Fig. 2m-n**) (gene sets too small for statistical functional enrichment evaluation).

### 4. Transcriptional response to sterilizing versus non-sterilizing antibiotics

Comparison of canonical sterilizing antibiotics (rifampin, pyrazinamide, bedaquiline) with non-sterilizing antibiotics (isoniazid, streptomycin, ethambutol) suggests that sterilizing drugs generate a more quiescent *Mtb* phenotype, as indicated by genes associated with translation, transcription, secretion of immunogenic proteins, metabolism, and cell wall synthesis. Specifically, expression of genes coding for primary ribosomal proteins, a basic metric of bacterial activity, was suppressed to a significantly greater degree by bedaquiline than by any non-sterilizing antibiotic (**Fig. 2a**). Rifampin and pyrazinamide suppressed primary ribosomal protein gene expression significantly more than two (isoniazid, ethambutol) of three non-sterilizing antibiotics. As discussed above, expression of the protein translation and modification gene category was decreased significantly for the sterilizing antibiotics but not for the non-sterilizing antibiotics. Expression of the gene for RNA polymerase subunit A (*rpoA*) was significantly decreased by all sterilizing antibiotics but not by any non-sterilizing antibiotics. Similarly, RNA polymerase subunit Z (*rpoZ)* was significantly decreased by all sterilizing antibiotics and only one (isoniazid) of the non-sterilizing antibiotics. All three sterilizing antibiotics had significantly decreased expression of *esxA,* the gene coding for ESAT-6, relative to isoniazid and ethambutol. Expression of the gene coding for isocitrate lyase (*icl1*), the first step of the glyoxylate bypass, was decreased significantly by all three sterilizing antibiotics but by none of the non-sterilizing antibiotics.

Expression of DosR regulon genes, which respond to hypoxia, carbon monoxide and nitric oxide encountered within host immune effector cells, was significantly reduced by all sterilizing drugs but not by the non-sterilizing drugs (**Fig. 3a**). Because bacterial DosR expression has previously been linked to the intensity of immune activation,^16,35^ we plotted the average scaled expression values for the ESX-1, ESX-3, and Antigen 85 genes against mean normalized expression of DosR regulon genes (**Fig. 3b-d**). Expression of ESX-1, ESX-3, and Antigen 85 were correlated with expression of the DosR regulon (R^2^=0.7, R^2^=0.745, R^2^=0.5, respectively), suggesting a link between bacterial phenotype and immune activation.

**Figure 3.**
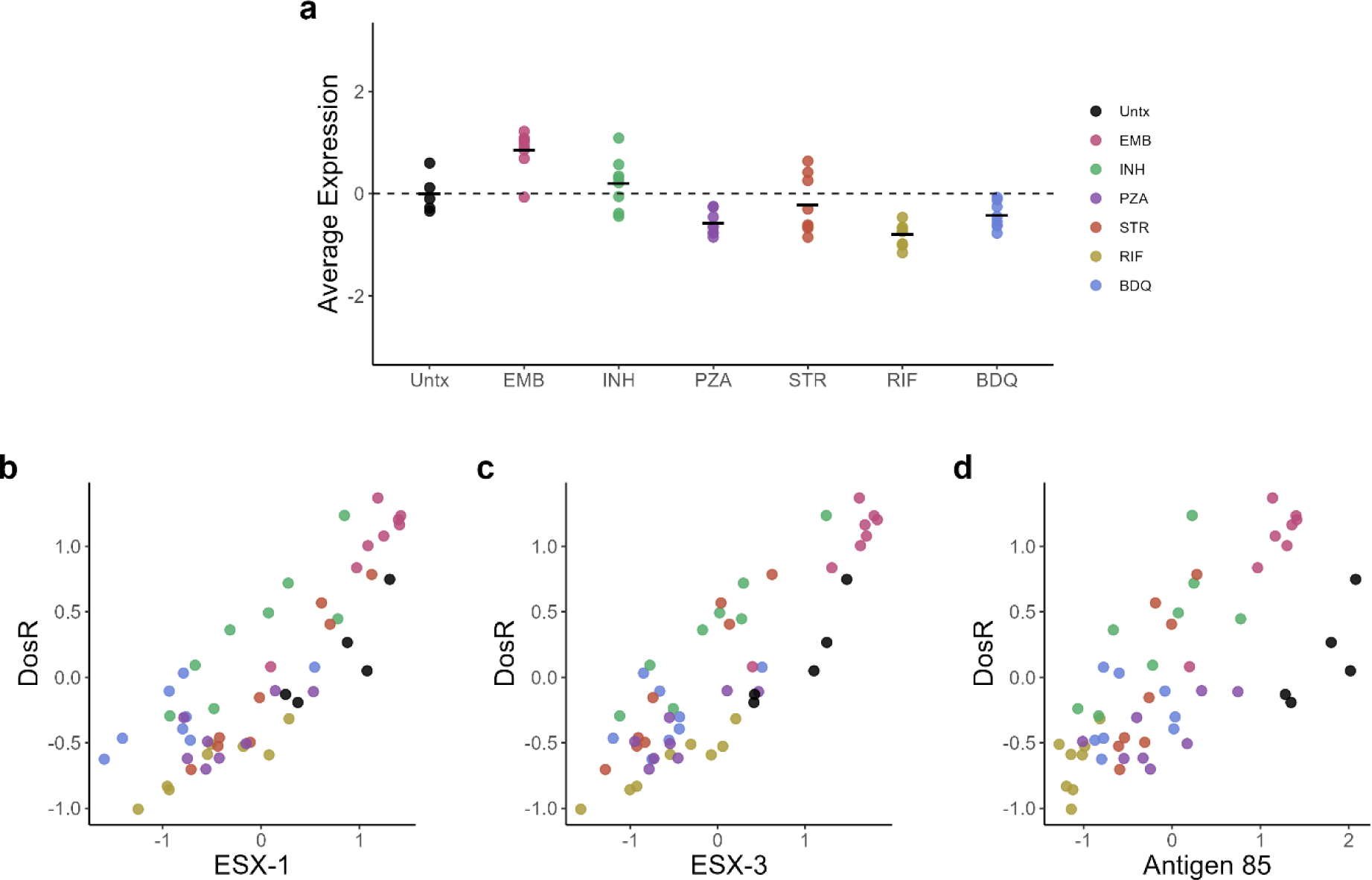
**a.** Average of VST-normalized, scaled expression across antibiotic treatments for genes in the DosR regulon. **b-c** Correlation between the scaled average expression for categories associated with immune activation and DosR: (**m)** ESX-1, (n) ESX-3, and (o) antigen 85. Each point represents an individual mouse and points are colored by treatment group. Abbreviations: Untreated (Untx), Ethambutol (EMB), Isoniazid (INH), Pyrazinamide (PZA), Streptomycin (STR), Rifampin (RIF), Bedaquiline (BDQ).

### 5. Distinguishing effects of individual antibiotics

Finally, we considered differences in transcriptional changes induced by each individual antibiotic exposure. Despite the existence of shared transcriptional changes discussed above, direct pairwise comparison between antibiotic exposures revealed that each antibiotic resulted in a distinct *Mtb* transcriptional response (**Fig. 4a**). Supplemental file 3 summarizes the categorical enrichment of each antibiotic to one another. Key observations from these tables are highlighted below.

**Figure 4.**
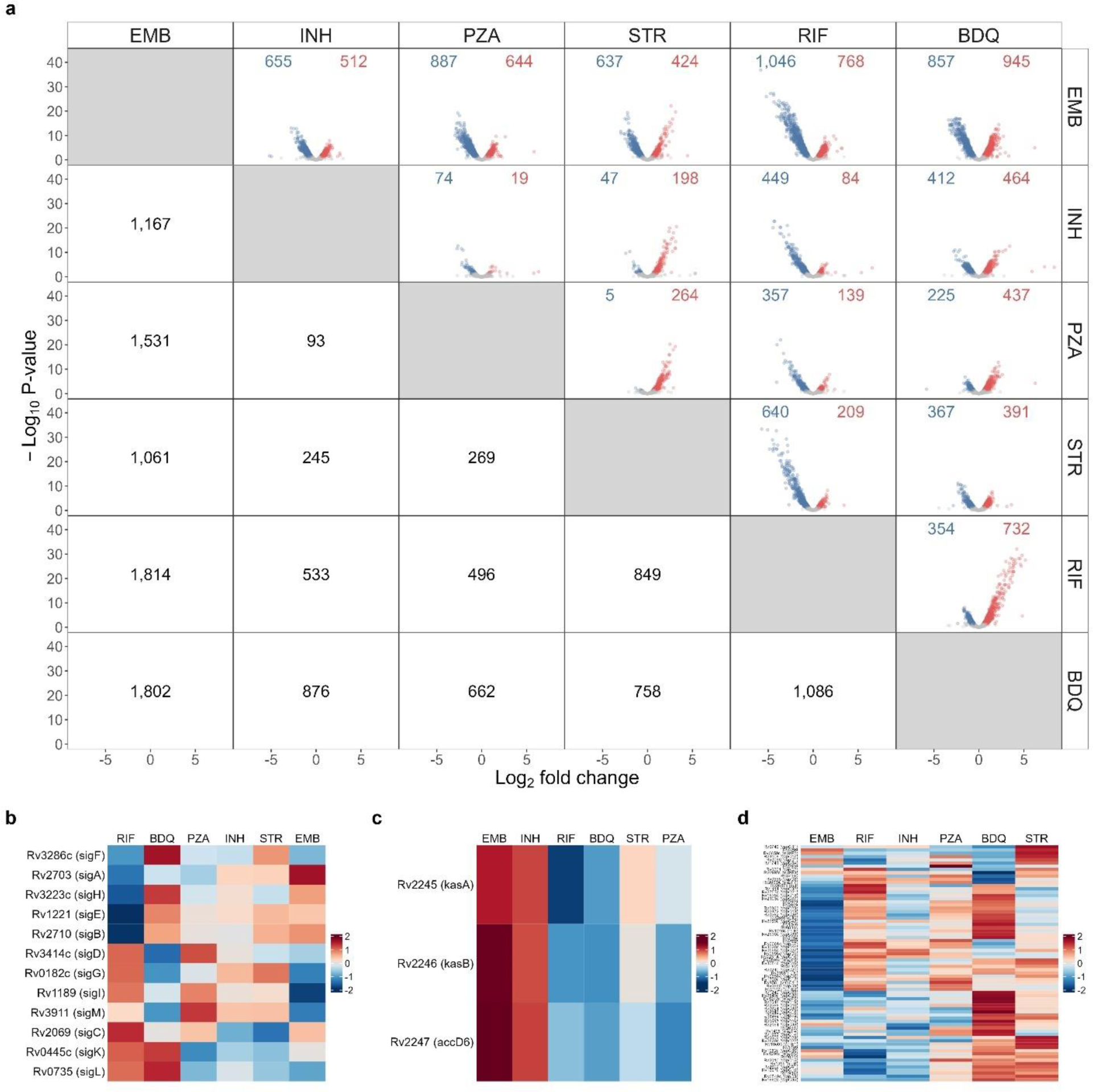
**a.** Differential expression in pairwise comparison between individual antibiotics. Volcano plots show fold change and significance between the antibiotics labeled in the row and column. The number of genes significantly down- (blue) or upregulated (red) with adj *p*-value< 0.05 in the row versus the column is shown below the diagonal. **b-d** Heatmaps showing the scaled average expression across antibiotic conditions for (**b**) sigma factors, (**c**) the *Kas* operon and (**d**) toxins. Abbreviations: Ethambutol (EMB), Isoniazid (INH), Pyrazinamide (PZA), Streptomycin (STR), Rifampin (RIF), Bedaquiline (BDQ).

#### Bedaquiline

Although evaluated at one-fifth the human-equivalent dose, bedaquiline induced the greatest transcriptional change of any antibiotic, significantly altering expression of 1,545 genes relative to untreated control (**Fig. 1c)**. The bedaquiline-treated phenotype was distinct, with at least 662 genes differentially expressed relative to any other antibiotic (bottom row of **Fig. 4a**). Inhibition of ATP synthetase via 4-week bedaquiline treatment led to a profoundly quiescent, inactive *Mtb* population, consistent with an energy-restricted phenotype. Specifically, relative to all antibiotics other than pyrazinamide, bedaquiline significantly decreased the expression of genes coding for primary ribosomal proteins and genes associated with the synthesis and modification of macromolecules. Bedaquiline suppressed the ESX1 locus to a significantly greater degree than isoniazid, streptomycin, or ethambutol. Additionally, bedaquiline induced greater expression of certain stress responses. Specifically, relative to any antibiotic other than streptomycin, bedaquiline induced significantly greater expression of genes for stressed-induced toxin/antitoxin modules. Relative to any other antibiotic, bedaquiline induced greater expression of sigma factor F, which directs growth arrest in response to diverse stresses (**Fig. 4b**).^36^

#### Rifampin

Evaluated at the existing standard human-equivalent dose, rifampin had the second-strongest effect on the *Mtb* transcriptome, significantly altering the expression of 1,000 genes relative to untreated control (**Fig. 1d**). The rifampin-treated phenotype was distinct, with at least 496 genes differentially expressed relative to any other antibiotic (second from bottom row of **Fig. 4a**). Rifampin resulted in significantly higher expression of genes involved in the cell wall than all antibiotics except ethambutol and significantly higher expression of PDIM than all antibiotics except ethambutol and isoniazid. Rifampin had significantly lower expression of the primary housekeeping sigma factor A than any antibiotic other than pyrazinamide, consistent with the regulation of a quiescent phenotype (**Fig. 4b** and Online Analysis Tool). Rifampin was distinct from all other antibiotics in having significantly increased expression of *sigE,* which codes for sigma factor E that mediates slower growth under stress conditions.^37^ All other antibiotics had significantly decreased expression of *sigE.* Rifampin resulted in significantly lower expression of genes coding for chaperones and heat shock proteins and the enduring hypoxic response^38^ than any other antibiotic. Rifampin-treated *Mtb* had significantly lower expression of the DosR regulon than *Mtb* treated with any antibiotic except bedaquiline.

#### Pyrazinamide

Pyrazinamide at human-equivalent dosing for 4 weeks resulted in broad changes in the *Mtb* transcriptome, significantly altering the expression of 822 genes relative to untreated control (**Fig. 1f)**. Because pyrazinamide had a static effect on CFU (no change relative to pre-treatment control, **Fig. 1a**), Pyrazinamide appears to induce adaptation of the existing *Mtb* population rather than selection of a pre-existing sub-population. Relative to rifampin, pyrazinamide had significantly higher expression of genes coding for the DosR regulon and the Antigen 85 complex as well as genes involved in beta-oxidation, electron transport, and toxin-antitoxin modules. Pyrazinamide clustered with isoniazid based on global similarity (**Fig. 1b,1i**) and relatively few genes were differentially expressed between pyrazinamide and isoniazid (96 significant genes, **Fig. 4a**), yet the pyrazinamide phenotype appeared less active than the isoniazid phenotype, with significantly lower expression of genes involved in protein translation and modification, ribosomal protein synthesis, and synthesis and modification of macromolecules.

#### Isoniazid

Isoniazid at human-equivalent dosing significantly altered the expression of 650 genes relative to untreated control (**Fig. 1g)**. Inhibition of mycolic acid synthesis by isoniazid was associated with higher expression of mycolic acid synthesis genes of the *kas* operon than any antibiotic other than ethambutol, suggesting continuing *Mtb* compensation to the isoniazid mechanism of action (**Fig. 4c**, Online Analysis Tool). Isoniazid also had significantly higher expression of DosR regulon genes compared to all antibiotics except ethambutol, suggesting adaptation to continued immune-mediated nitric oxide or hypoxic stress.

#### Streptomycin

Streptomycin at human-equivalent dosing significantly altered the expression of 850 genes relative to untreated control (**Fig 1e)**. The streptomycin phenotype was distinct, with at least 245 genes differentially expressed relative to any other antibiotic (**Fig. 4a**). Protein synthesis inhibition by streptomycin resulted in significantly higher expression of toxin-antitoxin pairs and of the enduring hypoxic response compared to any antibiotic other than bedaquiline. Streptomycin also resulted in significantly higher expression of chaperones and heat shock genes compared to any antibiotic other than ethambutol and significantly higher expression of genes associated with the response to oxidative stress than any antibiotic other than ethambutol or bedaquiline.

#### Ethambutol

Human-equivalent dosing of ethambutol induced the least transcriptional change among the antibiotics assessed, with 430 genes significantly altered relative to untreated control (**Fig. 1h**). The ethambutol transcriptome clustered with the untreated control (**Fig. 1i**), and was distinct from other antibiotics in most of the discrete processes shown in Figure 2.

## DISCUSSION

We found that 28-day treatment of mice with six different antibiotics led to emergence of antibiotic-specific *Mtb* transcriptional responses. Antibiotics differed both in the magnitude of transcriptional change they induced in *Mtb* and the specific sets of genes up- or down-regulated. Broadly, rifampin, pyrazinamide, and bedaquiline, the antibiotics with enhanced treatment-shortening activity (historically described as sterilizing), led to a less active bacterial phenotype than did antibiotics with lesser treatment-shortening activity (historically described as non-sterilizing).

*Mtb* phenotypes that lack resistance-conferring mutations, yet survive extended drug exposure *in vivo,* are viewed as a central obstacle to shortening the time required to cure TB.^39,40^ Our results suggest that different individual drugs result in distinct *in vivo* “persister” *Mtb* phenotypes. Rather, antibiotics with different mechanisms of action represent distinct injuries that condition the physiologic state of *Mtb* in distinct ways. While some broad transcriptional responses are shared among antibiotics (*e.g.,* down-regulation of genes associated with synthesis of macromolecules and metabolism and up-regulation of certain stress responses), each antibiotic also had unique effects on the *Mtb* transcriptome.

Of particular interest are sterilizing antibiotics known to play an outsized contribution to the ability of combination regimens to shorten the time required to TB cure. In this study, we selected three antibiotics with enhanced treatment-shortening activity – rifampin, pyrazinamide, and bedaquiline – that are central to contemporary regimen development and are included in recent and ongoing human trials. The SEARCH-TB analysis revealed that rifampin, pyrazinamide, and bedaquiline suppressed bacterial activity to a greater degree than did isoniazid, streptomycin, and ethambutol. This finding aligns with our previous observations using the RS ratio^®^ assay in the same mouse sample set which showed that rifampin, pyrazinamide, and bedaquiline decreased ribosomal RNA synthesis to a greater degree than antibiotics with lesser treatment-shortening activity.^41^ Combined with the RS ratio results, the SEARCH-TB data suggest that a common effect of antibiotics with potent treatment-shortening activity is the induction of a more inactive *Mtb* phenotype. Our findings suggest, but cannot definitively resolve, two potential interpretations for the observed association between treatment-shortening activity and decreased bacterial activity. First, a more quiescent phenotype may represent a functional physiologic adaptation that enables *Mtb* to survive exposure to rifampin, pyrazinamide, or bedaquiline, but is less crucial for surviving isoniazid, streptomycin, and ethambutol, streptomycin, and ethambutol. Alternatively, the more quiescent phenotype could be a “vital sign” of bacterial injury, signaling more severe stress and resultant bacterial dysfunction. *Mtb* population experiencing energy starvation (bedaquiline), or transcriptional inhibition (rifampin) may be functionally incapacitated or in a pre-terminal state.

For several antibiotics, the DosR regulon, which responds to nitric oxide and hypoxia *in vivo,* was downregulated relative to the untreated controls. This is consistent with previous observations in *Mtb* infected humans receiving antibiotic treatment^35^ and in mice treated with HRZE.^16^ Because antibiotics do not directly target generation of nitric oxide or restrict oxygen, the changes in the expression of the DosR regulon after antibiotic exposure is likely an indirect effect of treatment. Since activation of macrophages and neutrophils results in increased nitric oxide^42,43^ the observed downregulation in the DosR regulon after some antibiotic treatments may correspond to decreased inflammation. This theory is corroborated in the correlation of DosR regulon expression and the expression of the ESX-1 and ESX-3 systems, which have been linked with macrophage and neutrophil activation.^44,45^ If the observed fluctuation in the DosR regulon across antibiotic treatments is, in fact, a manifestation of host inflammation, this would indicate that antibiotics may impact host-pathogen interactions differently.

This work highlights the power of SEARCH-TB as a pharmacodynamic marker. In both preclinical studies and human trials, evaluation of new TB treatment has been hamstrung by limitations of existing culture-based pharmacodynamic markers.^46,47^ The fraction of the viable *Mtb* population that is capable of regrowth in culture is uncertain and may vary depending on antibiotic used.^48,49^ Additionally, enumeration of bacterial burden provides no information about how antibiotics affect *Mtb* physiologic processes. SEARCH-TB and other indicators of *Mtb* physiologic state such as the RS ratio reveal differences between drugs that appear identical based on burden. For example, we found that CFU did not distinguish between the effects of ethambutol (a weak antibiotic included in the standard regimen to protect against emergence of resistance) and pyrazinamide (an antibiotic shown to have potent treatment-shortening activity when added to combination regimens). By contrast, SEARCH-TB showed that pyrazinamide and ethambutol exposure resulted in profoundly different *Mtb* phenotypes with 1,531 genes or 43% of the transcriptome differentially expressed between the two. Similarly, isoniazid and rifampin, which are conventionally understood to play quite different roles in the existing standard regimen^18^, had indistinguishable effects on CFU but resulted in distinct molecular phenotypes. Our results indicate that antibiotics have effects that are not discernable based on the burden of bacilli recovered on solid agar. By evaluating bacterial physiologic processes rather than estimating bacterial burden, SEARCH-TB may reveal hitherto occult antibiotic effects that inform antibiotic development.

This report has several limitations. First, this report characterized drug-induced phenotypic change in the lungs of BALB/c mice, which develop loose macrophage aggregates containing intracellular *Mtb*. Other TB mouse models (such as the C3HeB/FeJ mouse) develop necrotic granulomas in which *Mtb* is extracellular and has distinct phenotypic adaptations to local conditions.^50^ A high-priority next step is interrogating *Mtb* in diverse models to elucidate the full spectrum of bacterial phenotypes and antibiotic responses. Second, we used the high-dose aerosol infection model because it is a mainstay of contemporary preclinical drug and regimen evaluation.^19^ High-dose aerosol infection is lethal if mice are not “rescued” by initiation of antibiotic treatment.^26^ In this experiment, untreated mice experienced clinical deterioration requiring humane euthanasia 19 days after aerosol infection. The untreated control therefore could not be temporally matched with the antibiotic-treated mice. Third, all antibiotics were evaluated at a human equivalent dose except bedaquiline which reduced *Mtb* burden below the limits of SEARCH-TB reliability at human equivalent dosing. It is likely that higher or lower drug doses might induce different transcriptional responses. Finally, SEARCH-TB quantifies expression in an entire lesion, inherently representing a population average that does not reveal heterogeneity within the population.

Using a novel pathogen-targeted RNA-seq method, we evaluated *Mtb* after 4-weeks of treatment with individual antibiotics *in vivo*, demonstrating that antibiotics with different mechanisms of action lead to distinct *Mtb* phenotypes. Sterilizing antibiotics generated a less active *Mtb* phenotype than non-sterilizing drugs. This report demonstrates the capability of SEARCH-TB to reveal differences in antibiotic effects that are not discernable via conventional microbiologic tools, potentially enabling a new era of pharmacodynamic monitoring in which candidate TB treatments are evaluated *in vivo* based on highly granular assessment of bacterial physiologic processes.

## REFERENCES

1. World Health Organization. Global tuberculosis report 2020. (2020).

2. Nahid, P. et al. Official American Thoracic Society/Centers for Disease Control and Prevention/Infectious Diseases Society of America clinical practice guidelines: Treatment of drug-susceptible tuberculosis. Clinical Infectious Diseases 63, e147–e195 (2016).

3. Dartois, V. A. & Rubin, E. J. Anti-tuberculosis treatment strategies and drug development: Challenges and priorities. Nature Reviews Microbiology 20, 685–701 (2022).

4. Roemhild, R., Bollenbach, T. & Andersson, D. I. The physiology and genetics of bacterial responses to antibiotic combinations. Nature Reviews Microbiology 20, 478–490 (2022).

5. Lenaerts, A., Barry, C. E. & Dartois, V. Heterogeneity in tuberculosis pathology, microenvironments and therapeutic responses. Immunol. Rev. 264, 288–307 (2015).

6. Sarathy, J. P. et al. Extreme drug tolerance of mycobacterium tuberculosis in caseum. Antimicrob. Agents Chemother. 62, (2018).

7. Lanni, F. et al. Adaptation to the intracellular environment of primary human macrophages influences drug susceptibility of *Mycobacterium tuberculosis*. Tuberculosis 139, 102318 (2023).

8. Abo-Kadoum, M. A., Dai, Y., Asaad, M., Hamdi, I. & Xie, J. Differential Isoniazid Response Pattern between Active and Dormant *Mycobacterium tuberculosis*. Microb. Drug Resist. 27, 768– 775 (2021).

9. Koul, A. et al. Delayed bactericidal response of *Mycobacterium tuberculosis* to bedaquiline involves remodelling of bacterial metabolism. Nat. Commun. 5, 3369 (2014).

10. Waddell, S. J. et al. The use of microarray analysis to determine the gene expression profiles of *Mycobacterium tuberculosis* in response to anti-bacterial compounds. Tuberculosis 84, 263–274 (2004).

11. Boshoff, H. I. M. et al. The transcriptional responses of *Mycobacterium tuberculosis* to inhibitors of metabolism: Novel insights into drug mechanisms of action. J. Biol. Chem. 279, 40174–40184 (2004).

12. Wilson, M. et al. Exploring drug-induced alterations in gene expression in *Mycobacterium tuberculosis* by microarray hybridization. Proc. Natl. Acad. Sci. U. S. A. 96, 12833–12838 (1999).

13. Deb, C. et al. A novel in vitro multiple-stress dormancy model for *Mycobacterium tuberculosis* generates a lipid-loaded, drug-tolerant, dormant pathogen. PLoS One 4, e6077 (2009).

14. Betts, J. C., Lukey, P. T., Robb, L. C., McAdam, R. A. & Duncan, K. Evaluation of a nutrient starvation model of *Mycobacterium tuberculosis* persistence by gene and protein expression profiling. Mol. Microbiol. 43, 717–731 (2002).

15. Poonawala, H. et al. Transcriptomic responses to antibiotic exposure in *Mycobacterium tuberculosis*. Antimicrob. Agents Chemother. 68, (2024).

16. Wynn, E. A. et al. Transcriptional adaptation of *Mycobacterium tuberculosis* that survives prolonged multi-drug treatment in mice. MBio (2023). doi:10.1128/mbio.02363-23

17. Mitchison, D. A. Basic mechanisms of chemotherapy. Chest 76, 771–780 (1979).

18. Mitchison, D. A. Role of individual drugs in the chemotherapy of tuberculosis. Int. J. Tuberc. Lung Dis. 4, 796–806 (2000).

19. Gumbo, T., Lenaerts, A. J., Hanna, D., Romero, K. & Nuermberger, E. Nonclinical models for antituberculosis drug development: A landscape analysis. J. Infect. Dis. 211, S83–S95 (2015).

20. Love, M. I., Huber, W. & Anders, S. Moderated estimation of fold change and dispersion for RNA-seq data with DESeq2. Genome Biol. 15, 550 (2014).

21. Robinson, M. D., McCarthy, D. J. & Smyth, G. K. edgeR: A Bioconductor package for differential expression analysis of digital gene expression data. Bioinformatics 26, 139–140 (2010).

22. Murtagh, F. & Legendre, P. Ward’s hierarchical agglomerative clustering method: Which algorithms implement Ward’s criterion? J. Classif. 31, 274–295 (2014).

23. Cole, S. T. et al. Erratum: Deciphering the biology of *Mycobacterium tuberculosis* from the complete genome sequence. Nature 396, 190 (1998).

24. Federico, A. & Monti, S. HypeR: An R package for geneset enrichment workflows. Bioinformatics 36, 1307–1308 (2020).

25. Benjamini, Y. & Hochberg, Y. Controlling the false discovery rate: A practical and powerful approach to multiple testing. J. R. Stat. Soc. Ser. B 57, 289–300 (1995).

26. Zhang, N. et al. Mechanistic modeling of *Mycobacterium tuberculosis* infection in murine models for drug and vaccine efficacy studies. Antimicrob. Agents Chemother. 64, (2020).

27. Prisic, S. et al. Zinc regulates a switch between primary and alternative S18 ribosomal proteins in *Mycobacterium tuberculosis*. Mol. Microbiol. 97, 263–280 (2015).

28. Kushwaha, A. K. & Bhushan, S. Unique structural features of the *Mycobacterium* ribosome. Progress in Biophysics and Molecular Biology 152, 15–24 (2020).

29. Karbalaei Zadeh Babaki, M., Soleimanpour, S. & Rezaee, S. A. Antigen 85 complex as a powerful *Mycobacterium tuberculosis* immunogene: Biology, immune-pathogenicity, applications in diagnosis, and vaccine design. Microbial Pathogenesis 112, 20–29 (2017).

30. Mascolo, L. & Bald, D. Cytochrome *bd* in Mycobacterium tuberculosis: A respiratory chain protein involved in the defense against antibacterials. Progress in Biophysics and Molecular Biology 152, 55–63 (2020).

31. Nandakumar, M., Nathan, C. & Rhee, K. Y. Isocitrate lyase mediates broad antibiotic tolerance in *Mycobacterium tuberculosis*. Nat. Commun. 5, (2014).

32. Garton, N. J. et al. Cytological and transcript analyses reveal fat and lazy persister-like bacilli in tuberculous sputum. PLoS Med. 5, 0634–0645 (2008).

33. Quigley, J. et al. The cell wall lipid PDIM contributes to phagosomal escape and host cell exit of *Mycobacterium tuberculosis*. MBio 8, (2017).

34. Klepp, L. I., Sabio y Garcia, J. & FabianaBigi. Mycobacterial MCE proteins as transporters that control lipid homeostasis of the cell wall. Tuberculosis 132, 102162 (2022).

35. Walter, N. D. et al. Transcriptional adaptation of drug-tolerant *Mycobacterium tuberculosis* during treatment of human tuberculosis. J. Infect. Dis. 212, 990–8 (2015).

36. Oh, Y. et al. The partner switching system of the SigF sigma factor in *Mycobacterium smegmatis* and induction of the SigF regulon under respiration-inhibitory conditions. Front. Microbiol. 11, 588487 (2020).

37. Chauhan, R. et al. Reconstruction and topological characterization of the sigma factor regulatory network of *Mycobacterium tuberculosis*. Nat. Commun. 7, (2016).

38. Rustad, T. R., Harrell, M. I., Liao, R. & Sherman, D. R. The enduring hypoxic response of *Mycobacterium tuberculosis*. PLoS One 3, e1502 (2008).

39. Connolly, L. E., Edelstein, P. H. & Ramakrishnan, L. Why is long-term therapy required to cure tuberculosis? PLoS Med. 4, e120 (2007).

40. Mandal, S., Njikan, S., Kumar, A., Early, J. V & Parish, T. The relevance of persisters in tuberculosis drug discovery. Microbiology (United Kingdom) 165, 492–499 (2019).

41. Walter, N. D., et al. *Mycobacterium tuberculosis* precursor rRNA as a measure of treatment-shortening activity of drugs and regimens. Nat. Commun. 12, 1–11 (2021).

42. Braverman, J. & Stanley, S. A. Nitric oxide modulates macrophage responses to *Mycobacterium tuberculosis* infection through activation of HIF-1α and Repression of NF-κB. J. Immunol. 199, 1805–1816 (2017).

43. Jamaati, H. et al. Nitric oxide in the pathogenesis and treatment of tuberculosis. Frontiers in Microbiology 8, 281243 (2017).

44. Conrad, W. H. et al. Mycobacterial ESX-1 secretion system mediates host cell lysis through bacterium contact-dependent gross membrane disruptions. Proc. Natl. Acad. Sci. U. S. A. 114, 1371–1376 (2017).

45. Tufariello, J. A. M. et al. Separable roles for *Mycobacterium tuberculosis* ESX-3 effectors in iron acquisition and virulence. Proc. Natl. Acad. Sci. U. S. A. 113, E348–E357 (2016).

46. Dooley, K. E., Phillips, P. P. J., Nahid, P. & Hoelscher, M. Challenges in the clinical assessment of novel tuberculosis drugs. Advanced Drug Delivery Reviews 102, 116–122 (2016).

47. Lenaerts, A. J., DeGroote, M. A. & Orme, I. M. Preclinical testing of new drugs for tuberculosis: Current challenges. Trends Microbiol. 16, 48–54 (2008).

48. Mukamolova, G. V., Turapov, O., Malkin, J., Woltmann, G. & Barer, M. R. Resuscitation-promoting factors reveal an occult population of tubercle bacilli in sputum. Am. J. Respir. Crit. Care Med. 181, 174–180 (2010).

49. Chengalroyen, M. D. et al. Detection and quantification of differentially culturable tubercle bacteria in sputum from patients with tuberculosis. Am. J. Respir. Crit. Care Med. 194, 1532–1540 (2016).

50. Walter, N. D. et al. Lung microenvironments harbor *Mycobacterium tuberculosis* phenotypes with distinct treatment responses. Antimicrob. Agents Chemother. 67, (2023).

