## Supplemental Information for "Emergence of antibiotic-specific *Mycobacterium tuberculosis* phenotypes during prolonged treatment of mice"

### 1 **Supplemental Methods**

#### 1.1 **Table S1.** Antibiotic dosing

**Table S1.** Antibiotic dosage concentration for mice in each antibiotic treatment group. Mice began treatment 11 days post infection via oral gavage five days a week for 28 days before euthanasia.

| **Treatment** | **Dose (mg/kg)** |
| --- | --- |
| Bedaquiline | 5 |
| Ethambutol | 100 |
| Isoniazid | 25 |
| Pyrazinamide | 150 |
| Rifampin | 10 |
| Streptomycin | 200 |

#### 1.3 **Figure S1.** Illustration of the method used to calculate and visualize the average expression of gene sets.

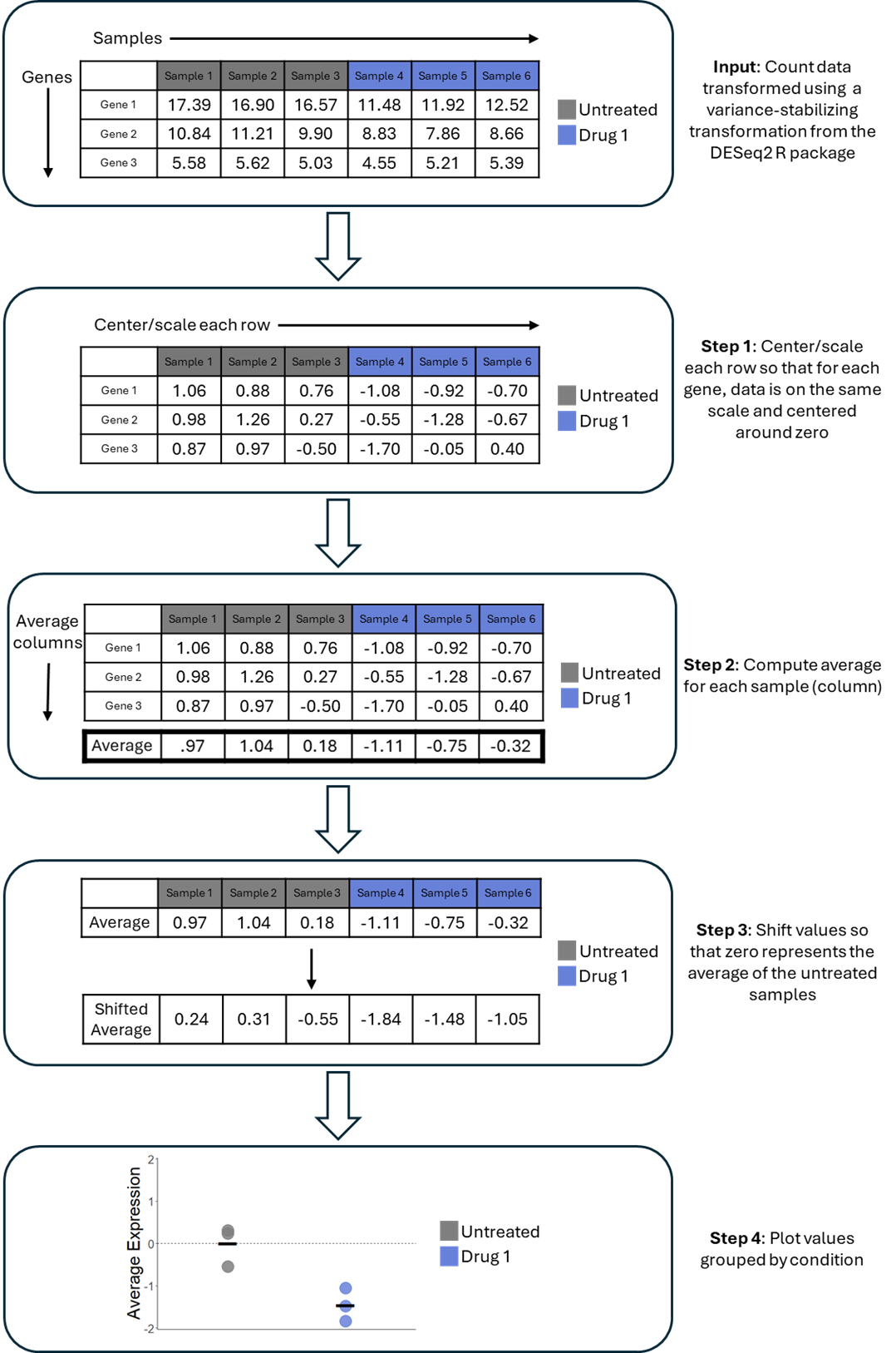

**Figure S1.** Illustration of the method for calculating and plotting the average expression of sets of genes. Mock data from two conditions (untreated and drug 1), each with three samples, is used for demonstration. For simplicity, a gene set of three genes is used. The variance-stabilized data for each gene is first scaled and centered around zero to ensure comparability. Then, the expression values are averaged for each sample and adjusted so that zero represents the average of the untreated samples. Finally, these values are plotted, providing a comparison of gene expression between conditions for the gene sets of interest and illustrating the spread of gene expression across samples.

#### 1.4 **Table S2.** Curated gene categories used for enrichment analysis.

**Table S2**. Manuscript figures and statistical functional enrichment analysis used sets of biologically linked genes curated from the literature. The number of genes in category indicates the number of genes identified in the literature source. Since SEARCH-TB quantifies expression of 89% of *Mtb* transcripts, not all genes in each category were quantified. Number of genes in SEARCH-TB indicates the number assayed.

| **Category** | **# of Genes in SEARCH-TB** | **# of Genes in Category** | **Source** |
| --- | --- | --- | --- |
| ABC transporters - Type I peptide and amino acids | 6 | 8 | (Soni, Dubey, & Bhatnagar, 2020)^1^ |
| ABC transporters - Type I Sugar Import | 12 | 12 | (Soni, Dubey, & Bhatnagar, 2020)^1^ |
| ABC transporters - Type I anion | 7 | 7 | (Soni, Dubey, & Bhatnagar, 2020)^1^ |
| ABC transporters - Type I phosphate | 8 | 8 | (Soni, Dubey, & Bhatnagar, 2020)^1^ |
| ABC transporters - Type II metal | 5 | 5 | (Soni, Dubey, & Bhatnagar, 2020)^1^ |
| Alternative ribosomal proteins | 4 | 5 | (Prisic et al., 2015)^2^ |
| Antigen 85 | 3 | 3 | (Karbalaei Zadeh Babaki, Soleimanpour, & Rezaee, 2017)^3^ |
| Antitoxins | 72 | 76 | (Shao et al., 2011)^4^ |
| Arabinogalactan (AG) | 18 | 19 | (Abrahams & Besra, 2018)^5^ |
| Beta Oxidation | 18 | 18 | (Schnappinger et al., 2003)^6^ |
| Cell wall synthesis | 40 | 40 | (Kirksey et al., 2011)^7^ |
| Cholesterol A and B ring degradation | 10 | 10 | (Pawełczyk et al., 2021)^8^ |
| Cholesterol C and D ring degradation | 5 | 5 | (Pawełczyk et al., 2021)^8^ |
| Cholesterol side chain degradation | 33 | 33 | (Pawełczyk et al., 2021)^8^ |
| Cutinase-Like Proteins (CULP) | 7 | 7 | (Tallman, Levine, & Beatty, 2016)^9^ |
| Cytochrome oxidase bccaa3 | 6 | 7 | (Lee, Sviriaeva, & Pethe, 2020) |
| Cytochrome oxidase bd | 3 | 4 | (Lee, Sviriaeva, & Pethe, 2020) |
| DNA replication and repair | 25 | 27 | (Ditse, Lamers, & Warner, 2017)^10^ |
| DosR | 48 | 48 | (Voskuil et al., 2003)^11^ |
| Drug targets | 40 | 40 | (“Working Group for New TB Drugs.,” 2021)^12^  (Shetye, Franzblau, & Cho, 2020)^13^ |
| Efflux Pumps and Transports | 25 | 26 | (Remm, Earp, Dick, Dartois, & Seeger, 2022)^14^ |
| Enduring Hypoxic Response | 149 | 161 | (Rustad, Harrell, Liao, & Sherman, 2008)^15^ |
| Esterases (Lip family) | 20 | 22 | (Tallman, Levine, & Beatty, 2016)^9^ |
| Esterases (non-Lip family) | 13 | 13 | (Tallman, Levine, & Beatty, 2016)^9^ |
| ESX1 | 18 | 19 | (Gröschel, Sayes, Simeone, Majlessi, & Brosch, 2016)^16^ |
| ESX2 | 12 | 12 | (Gröschel, Sayes, Simeone, Majlessi, & Brosch, 2016)^16^ |
| ESX3 | 9 | 11 | (Gröschel, Sayes, Simeone, Majlessi, & Brosch, 2016)^16^ |
| ESX4 | 7 | 7 | (Gröschel, Sayes, Simeone, Majlessi, & Brosch, 2016)^16^ |
| ESX5 | 11 | 15 | (Gröschel, Sayes, Simeone, Majlessi, & Brosch, 2016)^16^ |
| Fatty Acid Synthases I | 1 | 1 | (Cole et al., 1998)^17^ |
| Fatty Acid Synthases II | 8 | 9 | (Duan, Xiang, & Xie, 2014)^18^ |
| Fumarate reductase | 4 | 4 | (Cole et al., 1998)^17^ |
| Kas operon | 3 | 5 | (Slayden & Barry, 2002)^19^ |
| kstR1 regulon | 70 | 71 | (Wipperman, Sampson, & Thomas, 2014)^20^ |
| kstR2 regulon | 14 | 15 | (Wipperman, Sampson, & Thomas, 2014)^20^ |
| LpqY-SugA-SugB-Sug trehalose transporter | 5 | 5 | (Soni, Dubey, & Bhatnagar, 2020)^1^ |
| Mce1 | 7 | 7 | (Cole et al., 1998)^17^ |
| Mce2 | 6 | 7 | (Cole et al., 1998)^17^ |
| Mce3 | 7 | 7 | (Cole et al., 1998)^17^ |
| Mce4 | 7 | 7 | (Cole et al., 1998)^17^ |
| LAM | 14 | 15 | (Batt, Burke, Moorey, & Besra, 2020)^21^ |
| mmpL | 14 | 14 | (Domenech, Reed, & Barry, 2005)^22^ |
| mmpS | 5 | 5 | (Melly & Purdy, 2019)^23^ |
| Mycobactin Biogenesis | 10 | 10 | (Quadri, Sello, Keating, Weinreb, & Walsh, 1998)^24^ |
| Mycolic acid condensation and transfer | 7 | 7 | (Marrakchi, Lanéelle, & Daffé, 2014)^25^ |
| Mycolic acid modification | 12 | 12 | (Marrakchi, Lanéelle, & Daffé, 2014)^25^ |
| NADH dehydrogenase type I | 12 | 14 | (Cook, Hards, Vilchèze, Hartman, & Berney, 2014)^26^ |
| NADH dehydrogenase type II | 2 | 2 | (Cook, Hards, Vilchèze, Hartman, & Berney, 2014)^26^ |
| Nitrate import and reductase | 6 | 8 | (Cole et al., 1998)^17^ |
| Oxidative Stress | 48 | 49 | (Voskuil, Bartek, Visconti, & Schoolnik, 2011)^27^ |
| PDIM | 20 | 20 | (Rens, Chao, Sexton, Tocheva, & Av-Gay, 2021)^28^ |
| Peptidoglycan (PG) | 32 | 34 | (Maitra et al., 2019)^29^ |
| Phospholipase C | 4 | 4 | (Raynaud et al., 2002)^30^ |
| Primary ribosomal proteins | 50 | 53 | (Prisic et al., 2015)^2^, (Cole et al., 1998)^17^* |
| Sigma Factors | 12 | 13 | (Lew, Kapopoulou, Jones, & Cole, 2011)^31^ |
| Stringent Response - Induced | 58 | 70 | (Dahl et al., 2003)^32^ |
| Stringent Response - Repressed | 66 | 78 | (Dahl et al., 2003)^32^ |
| Succinate dehydrogenase I and II | 7 | 7 | (Hartman et al., 2014)^33^ |
| Toxin-Antitoxin | 146 | 152 | (Shao et al., 2011)^4^ |
| Toxins | 74 | 76 | (Shao et al., 2011)^4^ |
| Transcription Factors | 187 | 198 | (Lew, Kapopoulou, Jones, & Cole, 2011)^31^ |
| Trehalose | 10 | 10 | (Wilson et al., 1999)^34^ |
| Triacylglycerol Synthases | 14 | 16 | (Thanna & Sucheck, 2016)^35^ |
| UgpABCE glycerophosphocholine transporter | 4 | 4 | (Soni, Dubey, & Bhatnagar, 2020)1 |
| Universal stress proteins | 9 | 10 | (Lew, Kapopoulou, Jones, & Cole, 2011)^31^ |
| UspABC amino sugar transporter | 3 | 3 | (Soni, Dubey, & Bhatnagar, 2020)^1^ |
| WhiB-Like transcription factors | 7 | 7 | (Wan et al., 2021)^36^ |
| Zur regulon | 17 | 20 | (Dow et al., 2021)^37^ |

* Primary ribosomal proteins category was created by removing the four alternative proteins identified in Prisic et al., 2015 from the “Ribosomal protein and synthesis” gene list identified in Cole et al., 1998.

### 2 **Supplemental Results**

#### 2.1 **Figure S2:** Average expression for gene clusters from manuscript Figure 1i

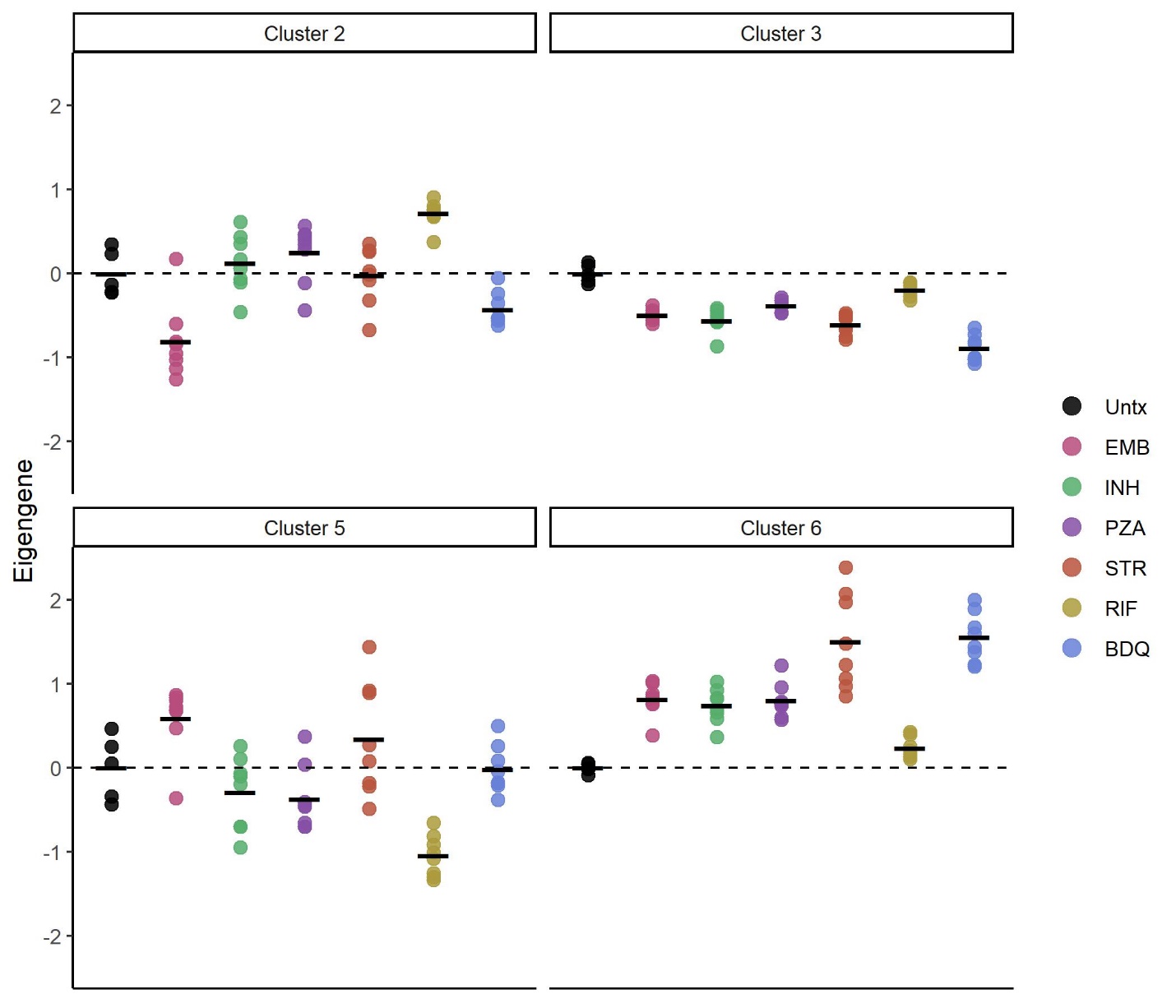

**Figure S2.** Scaled average gene expression of *Mycobacterium tuberculosis* in BALB/c infected mice across antibiotic treatments for clusters two, three, five and six from hierarchical gene clustering (Manuscript Fig. 1i). Each point represents an individual mouse. Horizontal lines indicate average values. Values are centered around the average value for the untreated samples so that points above and below zero represent upregulation or downregulation relative to untreated respectively. Abbreviations: Untreated (Untx), Ethambutol (EMB), Isoniazid (INH), Pyrazinamide (PZA), Streptomycin (STR), Rifampin (RIF), Bedaquiline (BDQ).

#### 2.2 **Figure S3**: Heatmaps of functional gene categories illustrated in manuscript Figure 2.

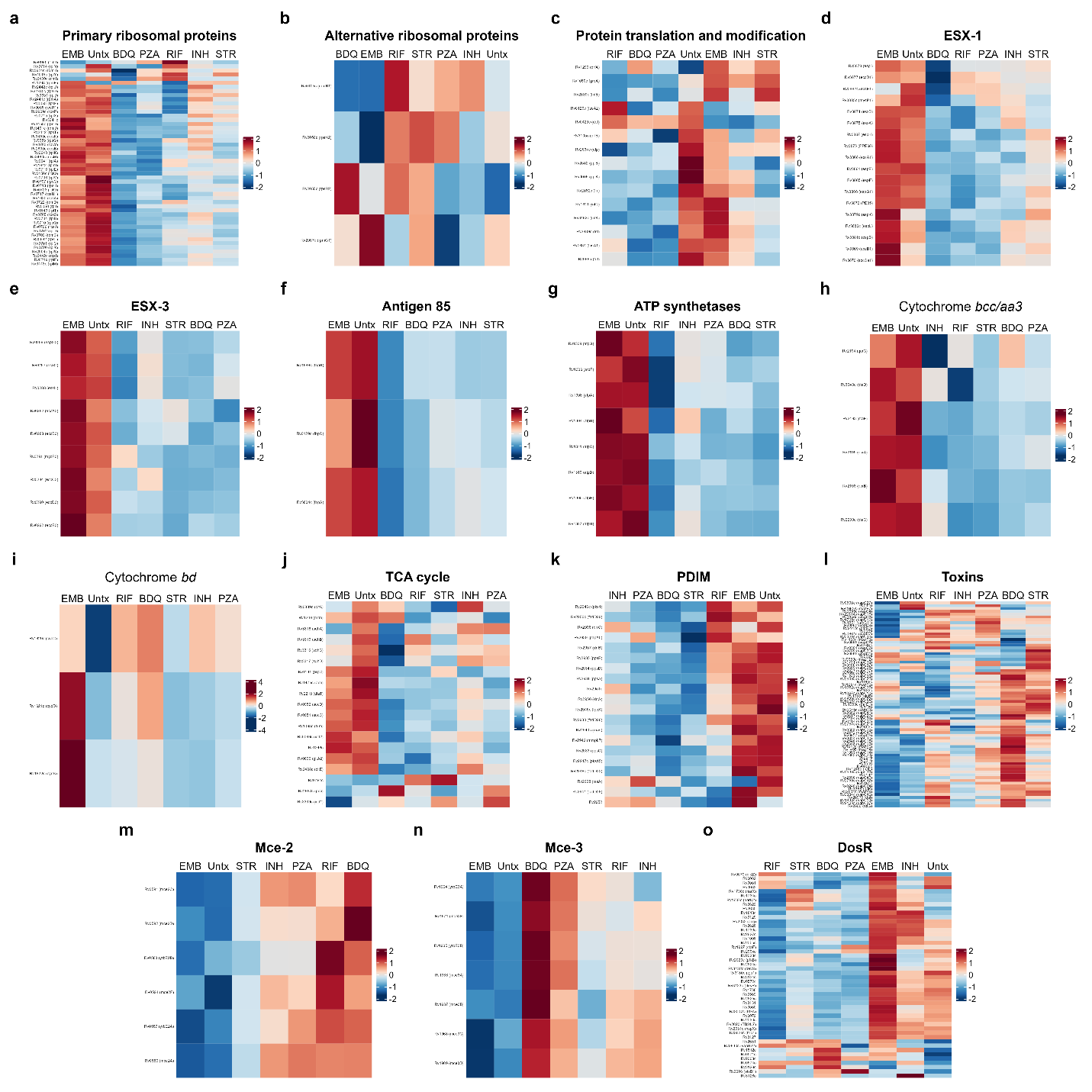

1.1 **Figure S3.** Heatmaps of functional gene categories illustrated in manuscript Figure 2.

**Figure S2.** Heatmaps showing the scaled average expression of *Mycobacteria tuberculosis* genes following antibiotic treatment in BALB/c mice for the functional gene categories shown in manuscript Figure 2. (**a**) primary ribosomal proteins, (**b**) alternative ribosomal proteins, (**c**) protein translation and modification, (**d**) ESX-1, (**e**) ESX-3, (**f**) antigen 85, (**g**) ATP synthesis, (**h**) cytochrome *bcc/aa3*, (**i**) cytochrome *bd*, (**j**) TCA cycle (**k**) PDIM, (**l**) toxins, (**m**) MCE-1, (**n**) MCE-3 and (**o**) DosR. Abbreviations: Untreated (Untx), Ethambutol (EMB), Isoniazid (INH), Pyrazinamide (PZA), Streptomycin (STR), Rifampin (RIF), Bedaquiline (BDQ).

### 2 **Supplemental references**

1.	Soni, D. K., Dubey, S. K. & Bhatnagar, R. ATP-binding cassette (ABC) import systems of *Mycobacterium tuberculosis*: target for drug and vaccine development. *Emerging Microbes and Infections* **9**, 207–220 (2020).

2.	Prisic, S. *et al.* Zinc regulates a switch between primary and alternative S18 ribosomal proteins in Mycobacterium tuberculosis. *Mol. Microbiol.* **97**, 263–280 (2015).

3.	Karbalaei Zadeh Babaki, M., Soleimanpour, S. & Rezaee, S. A. Antigen 85 complex as a powerful Mycobacterium tuberculosis immunogene: Biology, immune-pathogenicity, applications in diagnosis, and vaccine design. *Microbial Pathogenesis* **112**, 20–29 (2017).

4.	Shao, Y. *et al.* TADB: A web-based resource for Type 2 toxin-antitoxin loci in bacteria and archaea. *Nucleic Acids Res.* **39**, D606–D611 (2011).

5.	Abrahams, K. A. & Besra, G. S. Mycobacterial cell wall biosynthesis: A multifaceted antibiotic target. *Parasitology* **145**, 116–133 (2018).

6.	Schnappinger, D. *et al.* Transcriptional adaptation of *Mycobacterium tuberculosis* within macrophages: Insights into the phagosomal environment. *J. Exp. Med.* **198**, 693–704 (2003).

7.	Kirksey, M. A. *et al.* Spontaneous phthiocerol dimycocerosate-deficient variants of *Mycobacterium tuberculosis* are susceptible to gamma interferon-mediated immunity. *Infect. Immun.* **79**, 2829–2838 (2011).

8.	Pawełczyk, J. *et al.* Cholesterol-dependent transcriptome remodeling reveals new insight into the contribution of cholesterol to *Mycobacterium tuberculosis* pathogenesis. *Sci. Rep.* **11**, 1–16 (2021).

9.	Tallman, K. R., Levine, S. R. & Beatty, K. E. Small-molecule probes reveal esterases with Persistent Activity in dormant and reactivating *Mycobacterium tuberculosis*. *ACS Infect. Dis.* **2**, 936–944 (2016).

10.	Ditse, Z., Lamers, M. H. & Warner, D. F. DNA replication in *Mycobacterium tuberculosis*. *Microbiol. Spectr.* **5**, (2017).

11.	Voskuil, M. I. *et al.* Inhibition of respiration by nitric oxide induces a *Mycobacterium tuberculosis* dormancy program. *J. Exp. Med.* **198**, 705–713 (2003).

12.	Working Group for New TB Drugs. (2021). Available at: https://www.newtbdrugs.org/pipeline/drug-targets. (Accessed: 23rd February 2023)

13.	Shetye, G. S., Franzblau, S. G. & Cho, S. New tuberculosis drug targets, their inhibitors, and potential therapeutic impact. *Translational Research* **220**, 68–97 (2020).

14.	Remm, S., Earp, J. C., Dick, T., Dartois, V. & Seeger, M. A. Critical discussion on drug efflux in *Mycobacterium tuberculosis*. *FEMS Microbiology Reviews* **46**, 1–15 (2022).

15.	Rustad, T. R., Harrell, M. I., Liao, R. & Sherman, D. R. The enduring hypoxic response of *Mycobacterium tuberculosis*. *PLoS One* **3**, e1502 (2008).

16.	Gröschel, M. I., Sayes, F., Simeone, R., Majlessi, L. & Brosch, R. ESX secretion systems: Mycobacterial evolution to counter host immunity. *Nature Reviews Microbiology* **14**, 677–691 (2016).

17.	Cole, S. T. *et al.* Erratum: Deciphering the biology of Mycobacterium tuberculosis from the complete genome sequence. *Nature* **396**, 190 (1998).

18.	Duan, X., Xiang, X. & Xie, J. Crucial components of mycobacterium type II fatty acid biosynthesis (Fas-II) and their inhibitors. *FEMS Microbiol. Lett.* **360**, 87–99 (2014).

19.	Slayden, R. A. & Barry, C. E. The role of KasA and KasB in the biosynthesis of meromycolic acids and isoniazid resistance in Mycobacterium tuberculosis. *Tuberculosis* **82**, 149–160 (2002).

20.	Wipperman, M. F., Sampson, N. S. & Thomas, S. T. Pathogen roid rage: Cholesterol utilization by *Mycobacterium tuberculosis*. *Critical Reviews in Biochemistry and Molecular Biology* **49**, 269–293 (2014).

21.	Batt, S. M., Burke, C. E., Moorey, A. R. & Besra, G. S. Antibiotics and resistance: The two-sided coin of the mycobacterial cell wall. *Cell Surface* **6**, 100044 (2020).

22.	Domenech, P., Reed, M. B. & Barry, C. E. Contribution of the *Mycobacterium tuberculosis* MmpL protein family to virulence and drug resistance. *Infect. Immun.* **73**, 3492–3501 (2005).

23.	Melly, G. & Purdy, G. E. Mmpl proteins in physiology and pathogenesis of m. Tuberculosis. *Microorganisms* **7**, (2019).

24.	Quadri, L. E., Sello, J., Keating, T. A., Weinreb, P. H. & Walsh, C. T. Identification of a *Mycobacterium tuberculosis* gene cluster encoding the biosynthetic enzymes for assembly of the virulence-conferring siderophore mycobactin. *Chem. Biol.* **5**, 631–645 (1998).

25.	Marrakchi, H., Lanéelle, M. A. & Daffé, M. Mycolic acids: Structures, biosynthesis, and beyond. *Chemistry and Biology* **21**, 67–85 (2014).

26.	Cook, G. M., Hards, K., Vilchèze, C., Hartman, T. & Berney, M. Energetics of respiration and oxidative phosphorylation in mycobacteria. *Microbiol. Spectr.* **2**, (2014).

27.	Voskuil, M. I., Bartek, I. L., Visconti, K. & Schoolnik, G. K. The response of *Mycobacterium tuberculosis* to reactive oxygen and nitrogen species. *Front. Microbiol.* **2**, 105 (2011).

28.	Rens, C., Chao, J. D., Sexton, D. L., Tocheva, E. I. & Av-Gay, Y. Roles for phthiocerol dimycocerosate lipids in *Mycobacterium tuberculosis* pathogenesis. *Microbiology (United Kingdom)* **167**, 001042 (2021).

29.	Maitra, A. *et al.* Cell wall peptidoglycan in *Mycobacterium tuberculosis*: An Achilles’ heel for the TB-causing pathogen. *FEMS Microbiology Reviews* **43**, 548–575 (2019).

30.	Raynaud, C. *et al.* Phospholipases C are involved in the virulence of Mycobacterium tuberculosis. *Mol. Microbiol.* **45**, 203–217 (2002).

31.	Lew, J. M., Kapopoulou, A., Jones, L. M. & Cole, S. T. TubercuList - 10 years after. *Tuberculosis* **91**, 1–7 (2011).

32.	Dahl, J. L. *et al.* The role of RelMtb-mediated adaptation to stationary phase in long-term persistence of *Mycobacterium tuberculosis* in mice. *Proc. Natl. Acad. Sci. U. S. A.* **100**, 10026–10031 (2003).

33.	Hartman, T. *et al.* Succinate Dehydrogenase is the Regulator of Respiration in Mycobacterium tuberculosis. *PLoS Pathog.* **10**, (2014).

34.	Wilson, M. *et al.* Exploring drug-induced alterations in gene expression in *Mycobacterium tuberculosis* by microarray hybridization. *Proc. Natl. Acad. Sci. U. S. A.* **96**, 12833–12838 (1999).

35.	Thanna, S. & Sucheck, S. J. Targeting the trehalose utilization pathways of *Mycobacterium tuberculosis*. *MedChemComm* **7**, 69–85 (2016).

36.	Wan, T. *et al.* Structural insights into the functional divergence of WhiB-like proteins in Mycobacterium tuberculosis. *Mol. Cell* **81**, 2887-2900.e5 (2021).

37.	Dow, A. *et al.* Zinc limitation triggers anticipatory adaptations in *Mycobacterium tuberculosis*. *PLoS Pathog.* **17**, e1009570 (2021).
